# Antibody Evasion Properties of SARS-CoV-2 Omicron Sublineages

**DOI:** 10.1101/2022.02.07.479306

**Authors:** Sho Iketani, Lihong Liu, Yicheng Guo, Liyuan Liu, Yiming Huang, Maple Wang, Yang Luo, Jian Yu, Michael T. Yin, Magdalena E. Sobieszczyk, Yaoxing Huang, Harris H. Wang, Zizhang Sheng, David D. Ho

**Affiliations:** Aaron Diamond AIDS Research Center, Columbia University Vagelos College of Physicians and Surgeons, New York, NY 10032, USA; Department of Microbiology and Immunology, Columbia University Vagelos College of Physicians and Surgeons, New York, NY 10032, USA; Department of Systems Biology, Columbia University Vagelos College of Physicians and Surgeons, New York, NY 10032, USA; Division of Infectious Diseases, Department of Medicine, Columbia University Vagelos College of Physicians and Surgeons, New York, NY 10032, USA; Department of Pathology and Cell Biology, Columbia University Vagelos College of Physicians and Surgeons, New York, NY 10032, USA

**Author notes:** These authors contributed equally (Sho Iketani, Lihong Liu, Yicheng Guo, Liyuan Liu). Address correspondence to David D. Ho (, 701 W 168th St, 11th Floor, New York, NY 10032).

## Abstract

The identification of the Omicron variant (B.1.1.529.1 or BA.1) of SARS-CoV-2 (severe acute respiratory syndrome coronavirus 2) in Botswana in November 2021^1^ immediately raised alarms due to the sheer number of mutations in the spike glycoprotein that could lead to striking antibody evasion. We^2^ and others^3–6^ recently reported results in this Journal confirming such a concern. Continuing surveillance of Omicron evolution has since revealed the rise in prevalence of two sublineages, BA.1 with an R346K mutation (BA.1+R346K) and B.1.1.529.2 (BA.2), with the latter containing 8 unique spike mutations while lacking 13 spike mutations found in BA.1. We therefore extended our studies to include antigenic characterization of these new sublineages. Polyclonal sera from patients infected by wild-type SARS-CoV-2 or recipients of current mRNA vaccines showed a substantial loss in neutralizing activity against both BA.1+R346K and BA.2, with drops comparable to that already reported for BA.1^2,3,5,6^. These findings indicate that these three sublineages of Omicron are antigenically equidistant from the wild-type SARS-CoV-2 and thus similarly threaten the efficacies of current vaccines. BA.2 also exhibited marked resistance to 17 of 19 neutralizing monoclonal antibodies tested, including S309 (sotrovimab)^7^, which had retained appreciable activity against BA.1 and BA.1+R346K^2–4,6^. This new finding shows that no presently approved or authorized monoclonal antibody therapy could adequately cover all sublineages of the Omicron variant.

## Main Text

The meteoric rise of the B.1.1.529/Omicron to become the dominant SARS-CoV-2 variant globally has been truly remarkable^8^. Continuing surveillance of its evolution in the population over the past six weeks has revealed that the proportion of the original form, BA.1, has been decreasing steadily while the proportions of two other sublineages have increased noticeably (**Fig. 1a**). In fact, the BA.1+R346K sublineage now accounts for ∼30% of Omicron sequences globally, and ∼30-45% in South Africa, United Kingdom, and United States. On the other hand, the BA.2 sublineage accounts for only ∼13% of Omicron sequences globally, but it is not only on the rise but also the dominant form in countries such as Denmark and India. These three sublineages of Omicron share 21 mutations in the spike protein, wherein BA.2 contains 8 unique mutations and BA.1 contains 13 unique mutations (**Fig. 1b**). Of course, BA.1+R346K has one mutation more than BA.1. Given these differences, their antigenic properties cannot be assumed to be the same or similar.

**Fig. 1.**
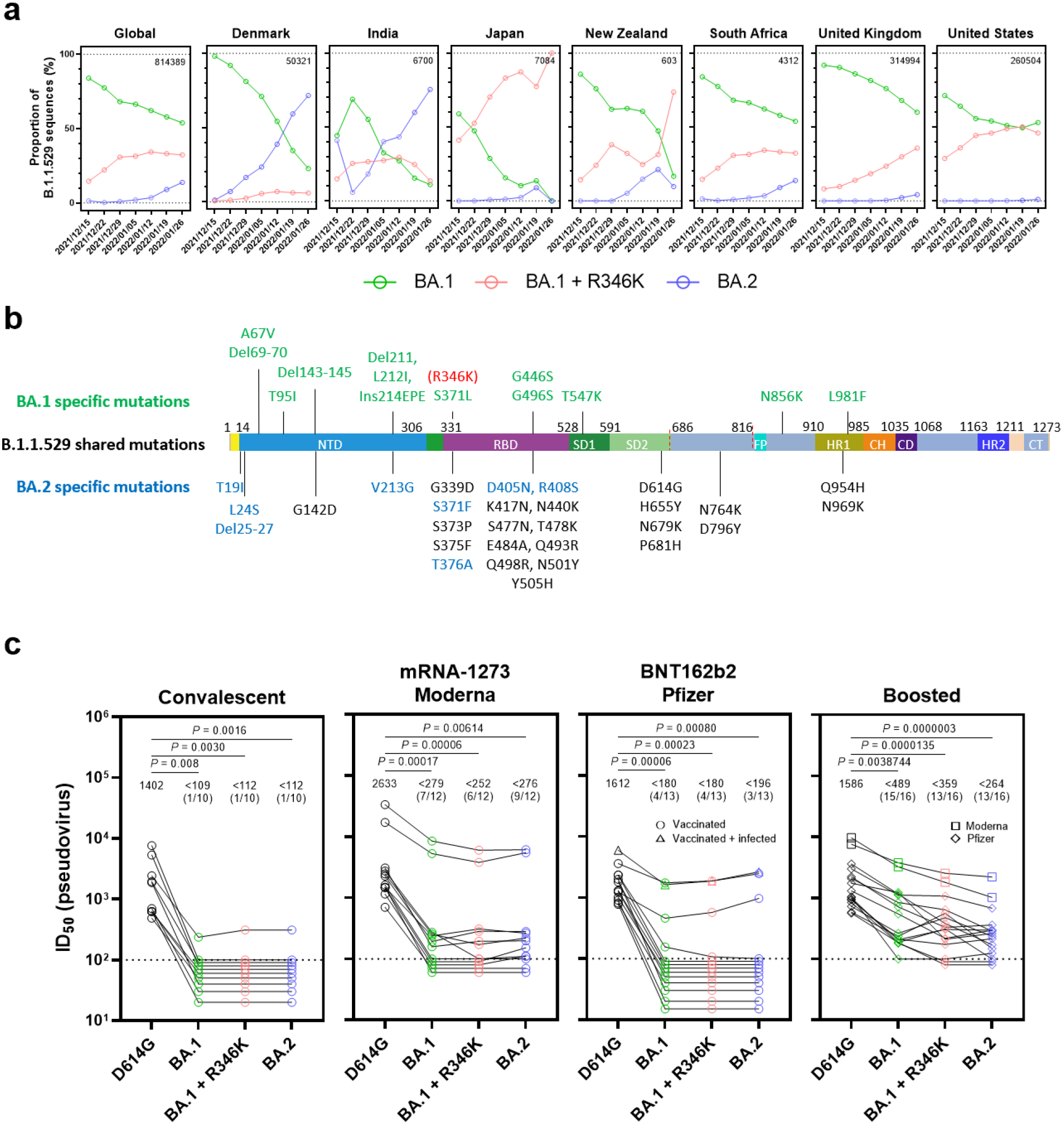
BA.2 exhibits a similar serum neutralization profile as BA.1 sublineages. **a,** Proportions of BA.1, BA.1+R346K, and BA.2 within B.1.1.529 sequences on GISAID over the past six weeks. Values in the upper right corner of each box denote cumulative number of Omicron sequences. **b,** Mutations within the B.1.1.529 lineage. **c,** Pseudovirus neutralization by convalescent and vaccinee sera. Values above points indicate the geometric mean. Numbers in parentheses denote the number of samples above the limit of detection (LOD) of 100. Values below the LOD are arbitrarily plotted to allow for visualization of each sample. P values were determined by two-sided Friedman test followed by Dunn’s multiple comparisons test.

Therefore, we first investigated the neutralization sensitivity of the Omicron sublineages by polyclonal sera from convalescent patients or individuals given mRNA vaccines, with or without a booster shot. These serum samples, as well as the pseudovirus neutralization assay used, were identical to ones previously reported^2^. The wild-type D614G pseudovirus was included as a comparator. As was observed and reported for BA.1^2,3,5,6^, a marked and significant loss of serum neutralizing activity against BA.1+R346K and BA.2 relative to D614G was noted, with neutralizing titers for numerous samples dropping below the limit of detection (**Fig. 1c**). The loss of neutralizing activity against BA.1+R346K or BA.2 sublineages was less prominent for sera obtained from individuals who received a booster vaccination (**Fig. 1c**, right panel), consistent with reported findings for BA.1^2,3,6^. Among these samples, the mean serum neutralizing titers against Omicron sublineages were significantly lower than the mean titer for D614G; although the mean titer was slightly lower for BA.2, the difference from BA.1 sublineages did not reach statistical significance (*P* = 0.242).

To further examine antigenic differences in the spike protein of these Omicron sublineages, a panel of 19 neutralizing monoclonal antibodies was used as probes. Seventeen were directed to different epitope clusters (classes 1-4) within the receptor-binding domain (RBD), whereas two were directed to the N-terminal domain (NTD). These antibodies included REGN10987 (imdevimab)^9^, REGN10933 (casirivimab)^9^, COV2-2196 (tixagevimab)^10^, COV2-2130 (cilgavimab)^10^, LY-CoV555 (bamlanivimab)^11^, CB6 (etesevimab)^12^, Brii-196 (amubarvimab)^13^, Brii-198 (romlusevimab)^13^, S309 (sotrovimab)^7^, LY-CoV1404 (bebtelovimab)^14^, ADG-2^15^, DH1047^16^, and S2X259^17^, as well as 1-20, 2-15, 2-7, 4-18, 5-7^18^ and 10-40^19^ from our group. Overall, 17 of 19 monoclonal antibodies were either totally inactive or severely impaired in neutralizing BA.2 (**Fig. 2a**), somewhat like previous findings for BA.1 and BA.1+R346K^2^ but with important differences (**Fig. 2b**). All class 4 antibodies tested lost greater neutralizing potency against BA.2 versus BA.1 sublineages. Two class 3 antibodies, COV2-2130 and 2-7, retained decent activity against BA.2 while having no activity against BA.1 viruses. S309 or sotrovimab lost 27-fold neutralizing activity against BA.2; this is particularly important because it was found to be the only clinically approved or authorized monoclonal antibody to retain activity against the original form of Omicron^2–4^. LY-CoV1404, another class 3 antibody in development, remained potent in neutralizing all Omicron sublineages, suggesting that there is still a patch within this antibody-binding region that is unaffected by all spike mutations found in SARS-CoV-2 variants to date. Although there was a lack of an observable difference among the Omicron sublineages in neutralization by polyclonal sera (**Fig. 1c**), important antigenic differences do exist when probed by monoclonal antibodies. Except for S309, BA.1 appears to be more resistant to class 3 antibodies than BA.2, while BA.2 is more resistant to all class 4 antibodies tested. Our recent study^2^ showed that previous SARS-CoV-2 variants, such as B.1.351/Beta and B.1.617.2/Delta, evolved to resist class 1, class 2, and NTD antibodies first, and then the Omicron variant seemingly has further evolved to resist class 3 and class 4 antibodies in addition. Our current findings suggest that the Omicron sublineages may have diverged under slightly different pressure from class 3 and class 4 antibodies to the RBD.

**Fig. 2.**
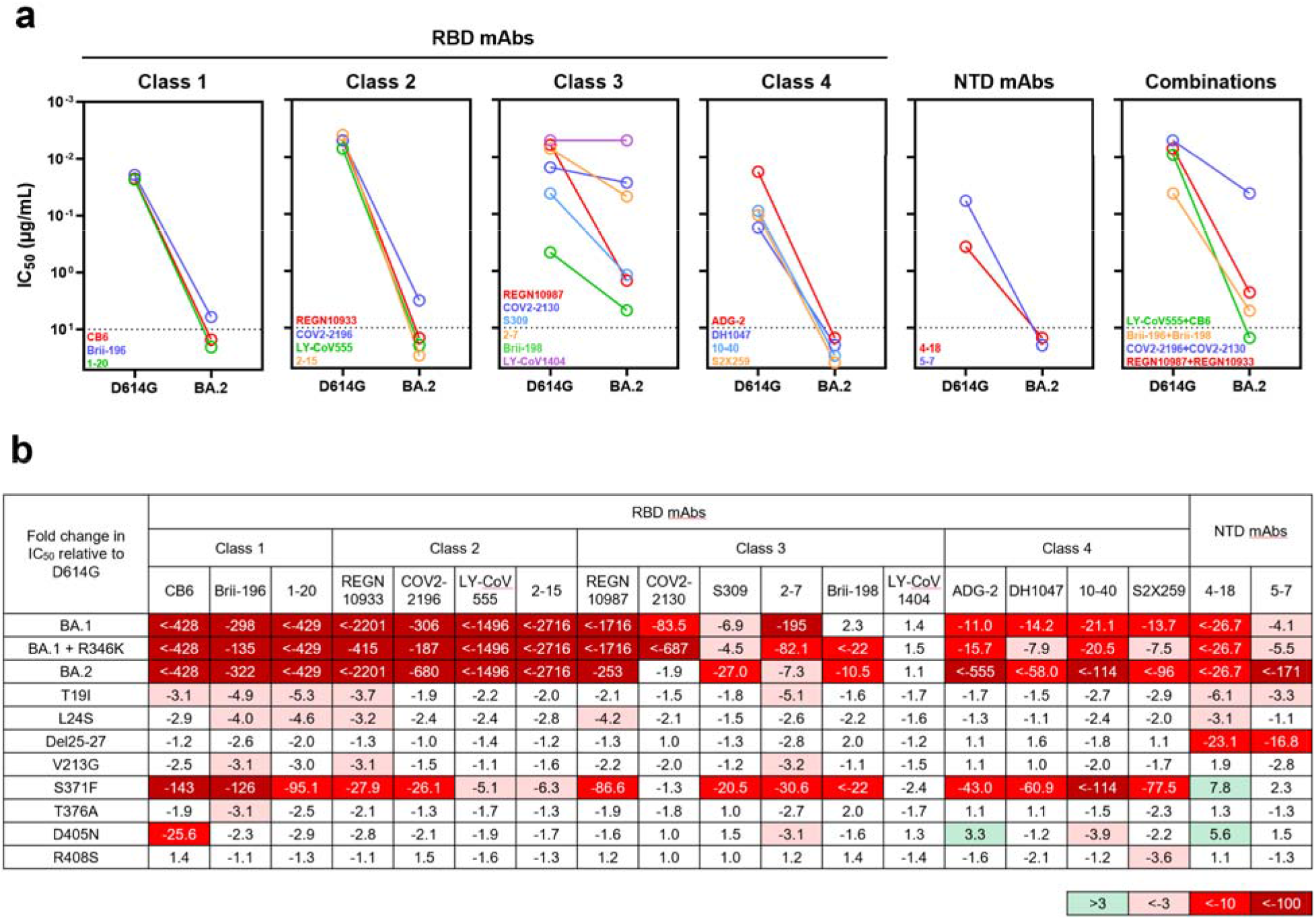
BA.2 differs in resistance profile to monoclonal antibodies. **a,** Pseudovirus neutralization by monoclonal antibodies. Values above the LOD of 10 μg/mL are arbitrarily plotted to allow for visualization of each sample**. b,** Fold change in IC_50_ values relative to D614G of neutralization of Omicron variants, as well as point mutants unique to BA.2.

Finally, we constructed each of the eight BA.2-specific spike mutations alone as pseudoviruses and tested them using the same panel of 19 monoclonal antibodies (**Fig. 2b**). S371F broadly affected most of the RBD-directed antibodies, similar to what was observed for S371L in BA.1^2^ but with a greater negative impact, perhaps due to the bulkier side chain of phenylalanine. Intriguingly but importantly, S371F appears to be majorly responsible for the loss in potency of S309, although this mutation was not observed previously as a marker for clinical resistance to sotrovimab^20^. CB6 was adversely affected by the D405N mutation, likely due to its position within the epitope of this antibody^12^. It is not clear how T19I and L24S mutations in the NTD subtly impaired the neutralizing activity of class 1 antibodies to RBD.

In summary, we have comprehensively evaluated the antigenic properties of two sublineages of the Omicron variant, BA.1+R346K and BA.2, and we believe our results have important clinical implications. First, polyclonal sera showed a substantial loss in neutralizing activity against both sublineages, with drops comparable to that of BA.1 (**Fig. 1c**). These three sublineages of Omicron, therefore, seem to be antigenically equidistant from the wild-type SARS-CoV-2, likely threatening the efficacies of current COVID-19 vaccines to a similar extent. The present study, however, does not address the antigenic distance between BA.1 and BA.2, which will require cross-neutralization experiments using sublineage-specific sera to determine. Second, monoclonal antibodies were affected in a disparate manner for the different Omicron sublineages. For clinically approved or authorized antibodies, only S309 (sotrovimab) retained activity against both BA.1 and BA.1+R346K, but its activity against BA.2 has dropped 27-fold (**Fig. 2b**) to a 50% inhibitory concentration (IC_50_) of ∼1 μg/mL (**Fig. 2a**). Only COV2-2130 (cilgavimab) and its combination with COV2-2196 (tixagevimab) retained activity against BA.2, but this antibody combination is only authorized for preventive use. Presently, no authorized therapeutic monoclonal antibody could adequately treat all sublineages of the Omicron variant. This finding poses a therapeutic dilemma in geographic regions where all three sublineages are present in sufficient numbers. As COVID-19 treatment options are narrowed by the emergence of more and more variants, it is imperative that we continue to devise novel strategies to contain this ever-evolving pathogen.

## Methods

### Data reporting

No statistical methods were used to predetermine sample size. The experiments were not randomized and the investigators were not blinded to allocation during experiments and outcome assessment.

### Serum samples

Identical samples from a previous study were utilized^2^. All collections were conducted under protocols reviewed and approved by the Institutional Review Board of Columbia University.

### Antibodies and pseudovirus neutralization

The expression of antibodies, construction of variant SARS-CoV-2 spike plasmids, production and neutralization of pseudoviruses, were conducted as previously described^2^.

## Acknowledgements

This study was supported by funding from the Gates Foundation, JPB Foundation, Andrew and Peggy Cherng, Samuel Yin, Carol Ludwig, David and Roger Wu, Regeneron Pharmaceuticals, the National Science Foundation (MCB-2032259), and the NIH SARS-CoV-2 Assessment of Viral Evolution (SAVE) Program.

## Author contributions

D.D.H. conceived this project. S.I. and Lihong Liu conducted pseudovirus neutralization experiments. Y.G. and Z.Z. conducted bioinformatic analyses. Liyuan Liu and Yiming Huang constructed the spike expression plasmids. M.W. aided sample collections. Y.L. managed the project. J.Y. expressed and purified antibodies. M.T.Y. and M.E.S. provided clinical samples. Yaoxing Huang contributed to discussions. H.H.W. and D.D.H. directed and supervised the project. S.I, Lihong Liu, and D.D.H. analyzed the results and wrote the manuscript.

## Data and materials availability

All data are provided in the manuscript.

